# Loss of allosteric regulation in α-isopropylmalate synthase identified as an antimicrobial resistance mechanism

**DOI:** 10.1101/2022.12.07.519492

**Authors:** Jaryd R. Sullivan, Christophe Courtine, Lorne Taylor, Ori Solomon, Marcel A. Behr

## Abstract

Despite our best efforts to discover new antimicrobials, bacteria have evolved mechanisms to become resistant. Resistance to antimicrobials can be attributed to innate, inducible, and acquired mechanisms. *Mycobacterium abscessus* is one of the most antimicrobial resistant bacteria and is known to cause chronic pulmonary infections within the cystic fibrosis community. Previously, we identified epetraborole as an inhibitor against *M. abscessus* with *in vitro* and *in vivo* activities and that the efficacy of epetraborole could be improved with the combination of the non-proteinogenic amino acid norvaline. Norvaline demonstrated activity against the *M. abscessus* epetraborole resistant mutants thus, limiting resistance to epetraborole in wild type populations. Here we show *M. abscessus* mutants with resistance to epetraborole can acquire resistance to norvaline in a leucyl-tRNA synthetase (LeuRS) editing-independent manner. After showing that the membrane hydrophobicity and efflux activity are not linked to norvaline resistance, whole-genome sequencing identified a mutation in the allosteric regulatory domain of α-isopropylmalate synthase (α-IPMS). We found that mutants with the α-IPMS^A555V^ variant incorporated less norvaline in the proteome and produced more leucine than the parental strain. Furthermore, we found that leucine can rescue growth inhibition from norvaline challenge in the parental strain. Our results demonstrate that *M. abscessus* can modulate its metabolism through mutations in an allosteric regulatory site to upregulate the biosynthesis of the natural LeuRS substrate and outcompete norvaline. These findings emphasize the antimicrobial resistant nature of *M. abscessus* and describe a unique mechanism of substrate-inhibitor competition.

**Significance Statement:** Cystic fibrosis patients and individuals undergoing plastic surgery are at risk for acquiring chronic infections from *Mycobacterium abscessus*. Current antibiotics are not adequate and require increased drug discovery efforts to identify better treatments for these patients. The benzoxaborole, epetraborole has been shown by our group and others to be a promising candidate against *M. abscessus* but the emergence of resistance to epetraborole in a clinical trial for complicated urinary tract infections has hindered its development. Previously, we identified the combination of epetraborole and norvaline as a potential means to limit resistance against epetraborole. Our results here demonstrate that *M. abscessus* can acquire resistance to both epetraborole and norvaline. These results may help develop combination therapies to reduce the risk of resistance to benzoxaboroles and non-proteinogenic amino acids.

## Introduction

Antimicrobial resistance (AMR) is a major public health concern and is estimated to contribute to 5 million deaths each year (1–3). To facilitate antimicrobial research and development, the WHO created a priority pathogen list based on trends of resistance, preventability, and the status of the drug pipeline (4, 5). Pathogens on this list are considered critical, high, or medium priority with a separate designation for *Mycobacterium tuberculosis*. Arguably, when compared to *M. tuberculosis, M. abscessus* is more drug resistant, more recalcitrant to treatment, and has less lead compounds in its pipeline than *M. tuberculosis*, yet it is often neglected in discussions regarding the current threat of antimicrobial resistant bacteria (6–10).

Resistance to antimicrobials can be attributed to innate, inducible, and acquired mechanisms. *M. abscessus* uses a variety of mechanisms to overcome the antibiotic regimens clinicians have at their disposal. Besides acquired target mutations that alter the binding affinity for antibiotics like clarithromycin (11), *M. abscessus* has a complex phenotypic resistome. The *M. abscessus*/*M. chelonae* complex is recognized for its unusually hydrophobic membrane relative to *M. tuberculosis*, which decreases the permeability to small nutrients and amino acids, and antibiotics across the cell wall (12, 13). Moreover, it has also been established that the cell wall synergizes with the internal resistome that includes efflux pumps and drug modifying enzymes to limit the effect of toxic compounds (6). *M. abscessus* expresses the ADP-ribosyltransferase *arr*_Mab_ that ribosylates rifamycins, and the *N*-acetyltransferase *eis2* and 3’-O-phosphotransferase *aph(3’)* that inactivate aminoglycosides (14–17). The genome also encodes the β-lactamases Bla_Mab_, which can hydrolyze penicillins and cephalosporins, and Bla_Mmas_, which has extended spectrum activity against carbapenems to limit the availability of the most widely-used antibiotic class used to treat bacterial infections (18–20). Many of the mechanisms listed here are under the regulation of inducible transcription factors like WhiB7. WhiB7 is known to be activated by ribosomal protein synthesis inhibitors like macrolides, tetracyclines, and aminoglycosides but also by molecules with pleiotropic effects on cell metabolism (21–23). It was shown in *M. abscessus* that inducible macrolide resistance is mediated through upregulation of *erm*(41) in a WhiB7-dependent process (24–26). These mechanisms act in synergy with a hydrophobic membrane composed of mycolic acids with significantly less permeability than the membranes of other mycobacteria, resulting a majority of treatment failures (12, 13, 27, 28).

By elucidating AMR mechanisms and leveraging the bacterial response to antimicrobial agents, novel antimicrobials and combination therapies could be developed for *M. abscessus* infections. Previously, we identified the leucyl-tRNA synthetase (LeuRS) inhibitor epetraborole with nanomolar whole cell activity against *M. abscessus* (29). During this screening program, we discovered that epetraborole had failed a phase 2 clinical trial due to the rapid emergence of resistant *Escherichia coli* in complicated urinary tract infections (30). We showed that epetraborole resistant *M. abscessus* mutants became sensitive to the non-proteinogenic amino acid l-norvaline. Sensitivity to l-norvaline is derived from a D436H substitution in the editing domain of LeuRS, which is the binding site for the epetraborole-tRNA^Leu^ adduct. LeuRS activates l-leucine and subsequently transfers the activated amino acid onto the corresponding tRNA^Leu^ isoacceptors. Fortunately, tRNA^Leu^ with non-cognate amino acids can undergo hydrolysis in the editing domain to remove the erroneous amino acid before participating in polypeptide synthesis. However, by gaining resistance to epetraborole from LeuRS^D436H^, the ability to edit misaminoacylated tRNA^Leu^ is lost and the risk of translating incorrect proteins increases. Treatment of the epetraborole resistant mutants with l-norvaline caused proteome-wide misincorporation with l-norvaline at residues coding for l-leucine, and a decrease in the emergence of resistance to epetraborole when given in combination.

In this article, we show using a combination of genetics, proteomics, and small molecule analysis that overproduction of l-leucine from insensitivity to feedback inhibition can abrogate the toxicity of l-norvaline to levels similar to using the evolutionarily conserved editing domain on LeuRS. These findings demonstrate the loss of allosteric regulation as the foundation for competitive antimicrobial inhibition and highlight the propensity of *M. abscessus* to become increasingly antimicrobial resistant.

## Results

### Spontaneous Generation of l-Norvaline Resistance in *M. abscessus*

*M. abscessus* that is resistant to both epetraborole and l-norvaline was raised spontaneously during a kill kinetics experiment that sought to determine whether l-norvaline caused cell death to the epetraborole resistant *M. abscessus* LeuRS^D436H^ strain. The wild type ATCC 19977 reference strain of *M. abscessus*, the EPT^R^ LeuRS^D436H^ mutant, and the EPT^R^ LeuRS^D436H^ mutant complemented with wild type *leuS* were grown in 2.4 mM l-norvaline (4X MIC) for five days. In agreement with the editing activity of LeuRS, the wild type and complemented strain grew in 2.4 mM l-norvaline, unlike the editing-deficient mutant (Fig. 1*A*). For the latter strain, l-norvaline challenge resulted in a loss of viable bacteria, congruent with the toxicity from misfolded proteins. Unexpectedly, the growth of EPT^R^ LeuRS^D436H^ mutant recovered at later timepoints. We hypothesized that l-norvaline might degrade over time and repeated the experiment where 2.4 mM l-norvaline was supplemented each day to create a steady-state (4XSS). Again, however, the EPT^R^ LeuRS^D436H^ mutant grew after day three (Fig. 1*A*). To confirm resistance to l-norvaline, aliquots of the EPT^R^ LeuRS^D436H^ mutant grown without l-norvaline and in 4XSS were taken to measure the MIC_90_ of l-norvaline using the resazurin microtiter assay (REMA) method. Indeed, the EPT^R^ LeuRS^D436H^ 4XSS strain had a MIC_90_ > 40 mM while the original EPT^R^ LeuRS^D436H^ strain grown without l-norvaline had a MIC_90_ of 0.2 mM.

**Figure 1.**
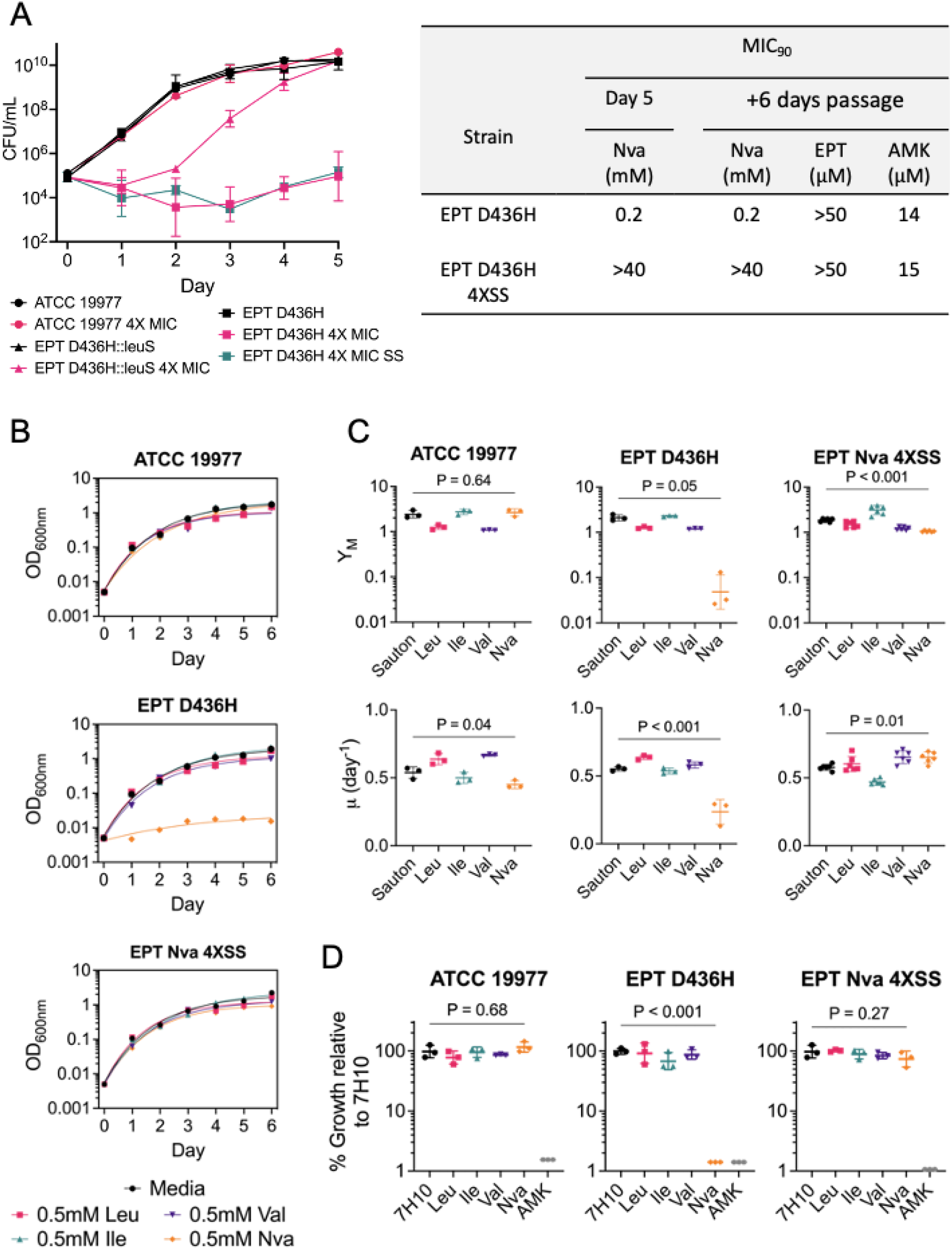
Editing-deficient norvaline resistance in *M. abscessus*. (A) Left, WT *M. abscessus* ATCC 19977, EPT^R^ *M. abscessus* with LeuRS^D436H^, and complement strain with WT LeuRS were grown in media free of exogenous l-norvaline, 4X MIC_90_ (MIC_90_ = 0.6 mM) of the Nva^S^ /EPT^R^ *M. abscessus* D436H strain. Alternatively, *M. abscessus* D436H strains were grown in media free of exogenous l-norvaline, with an initial inoculum of 4X MIC_90_, or media replenished with 4X MIC_90_ daily to establish a steady-state of l-norvaline (SS). Data represents mean ± SD from n = 2-5 independent experiments. Right, Aliquots of *M. abscessus* EPT^R^ D436H grown in l-norvaline free media (black squares) or l-norvaline at 4XSS (teal squares) from day 5 were used to determine the MIC_90_ of l-norvaline. Cultures were passaged for six days in l-norvaline free media before measuring the MIC_90_ for l-norvaline, and control drugs epetraborole and amikacin. (B) Growth curves of *M. abscessus* ATCC 19977, *M. abscessus* EPT^R^ D436H, and mutant with dual EPT^R^ and Nva^R^ resistance raised at 4XSS. Strains were grown statically in 96-well plates with nutrient limited Sauton media ± BCAAs and fit with exponential plateau regression. Data is representative of n = 3 independent experiments with mean ± SD. P values were obtained by one-way ANOVA with Dunnett’s multiple comparisons test. (C) Comparison of maximum growth plateau (Y_M_) and growth rates (μ) in Sauton media ± BCAAs. Data is mean ± SD from n = 3 independent replicates with *M. abscessus* EPT^R^ Nva^R^ 4XSS in biological duplicates. (D) Growth on solid 7H10 media ± BCAAs. 100 μM amikacin was used as control. Data is mean ± SD from n = 3 independent experiments. P values were obtained by one-way ANOVA with Dunnett’s multiple comparisons test.

Since *M. abscessus* was shown to have inducible resistance to macrolides (31, 32), we asked whether l-norvaline resistance was inducible or acquired. We previously showed that inducible macrolide resistance in *M. abscessus* can be repressed after passaging the culture in drug-free media for six days (29). Aliquots of the EPT^R^ LeuRS^D436H^ grown in l-norvaline-free or 4XSS conditions on day five were passaged for six days in l-norvaline-free media. The MIC_90_ to l-norvaline, epetraborole, and amikacin were determined using REMA. We showed that l-norvaline resistance is not inducible since passaging the culture in l-norvaline-free media did not change the MIC_90_ (Fig. 1*A*). Alternatively, l-norvaline resistance could arise from a LeuRS^D436H^ reversion where the population regains editing activity but becomes susceptible to epetraborole. Refuting this possibility, resistance to epetraborole was retained with MIC_90_ > 0.05 mM (Fig. 1*A*). Lastly, l-norvaline resistance was not the result of a cross-resistance as evidenced with sensitivity to the control drug amikacin.

To further characterize the new mutant resistant to both epetraborole and l-norvaline, we challenged *M. abscessus* ATCC 19977, *M. abscessus* EPT^R^ LeuRS^D436H^, and *M. abscessus* EPT^R^ Nva^R^ 4XSS with 0.5 mM l-norvaline and other branched-chain amino acids (BCAAs) in Sauton’s minimal media and fit their growth to an exponential plateau regression (Fig. 1*B*). As controls, *M. abscessus* ATCC 19977 grew in l-norvaline while *M. abscessus* EPT^R^ LeuRS^D436H^ failed to grow. Like the wild type strain, the growth of *M. abscessus* EPT^R^ Nva^R^ 4XSS was not impeded by 0.5 mM l-norvaline. From the exponential fit, we extracted the maximum growth plateau (Y_M_) and growth rate (μ). Quantitatively, we observed that l-norvaline impairs both the maximum growth and growth rate of an editing deficient strain like *M. abscessus* EPT^R^ LeuRS^D436H^ but that *M. abscessus* EPT^R^ Nva^R^ 4XSS regained a normal growth rate with a lower plateau (Fig. 1*C*). Similar results were obtained when the experiments were repeated in nutrient rich 7H9 media (Fig. S1).

Growth in 0.5 mM BCAAs was repeated on solid media using 7H10 base and 100 μM amikacin as control. We observed similar results where only *M. abscessus* EPT^R^ LeuRS^D436H^ failed to grow on 0.5 mM l-norvaline while amikacin inhibited all strains (Fig. 1*D*). These results (*i*) confirmed l-norvaline resistance and (*ii*) suggested a novel mechanism of l-norvaline resistance acquired by *M. abscessus* that is editing independent.

### Membrane Hydrophobicity does not Drive Norvaline Resistance

To examine whether the *M. abscessus* EPT^R^ Nva^R^ 4XSS mutant has a modified cell envelope, we measured the hydrophobicity via the uptake of Congo red dye. Strains were streaked onto 7H10 plates with 10% (v/v) OADC and 140 μM Congo red and incubated at 37 °C for five days (Fig. 2*A*). *M. abscessus* ATCC 19977 strains with the smooth morphotype (S) or rough morphotype (R) were used as representative hydrophilic and hydrophobic membranes, respectively. The rough morphotype contains a mutation in *mmpL4b*, which prevents the transport of glycopeptidolipids across the membrane and creates a uniform hydrophobicity across the cell envelope unlike the smooth morphotype that has clusters of hydrophobic domains (33). Gross visualization of the colonies indicated that both the parental EPT^R^ LeuRS^D436H^ and EPT^R^ Nva^R^ 4XSS strains had a smooth morphotype which favors pink colonies with red borders while the rough morphotype colonies were red (Fig. 2*A*). Congo red dye retained in the membrane was extracted with DMSO and quantified at 488nm. We observed a significant increase in retained Congo red dye in *M. abscessus* R compared to S, however there was no significant change in dye retained in EPT^R^ Nva^R^ 4XSS relative to the parental strain EPT^R^ LeuRS^D436H^ (Fig. 2*B*). These results suggest that the cell membrane of *M. abscessus* EPT^R^ Nva^R^ 4XSS has not undergone modifications to decrease the permeability to l-norvaline.

**Figure 2.**
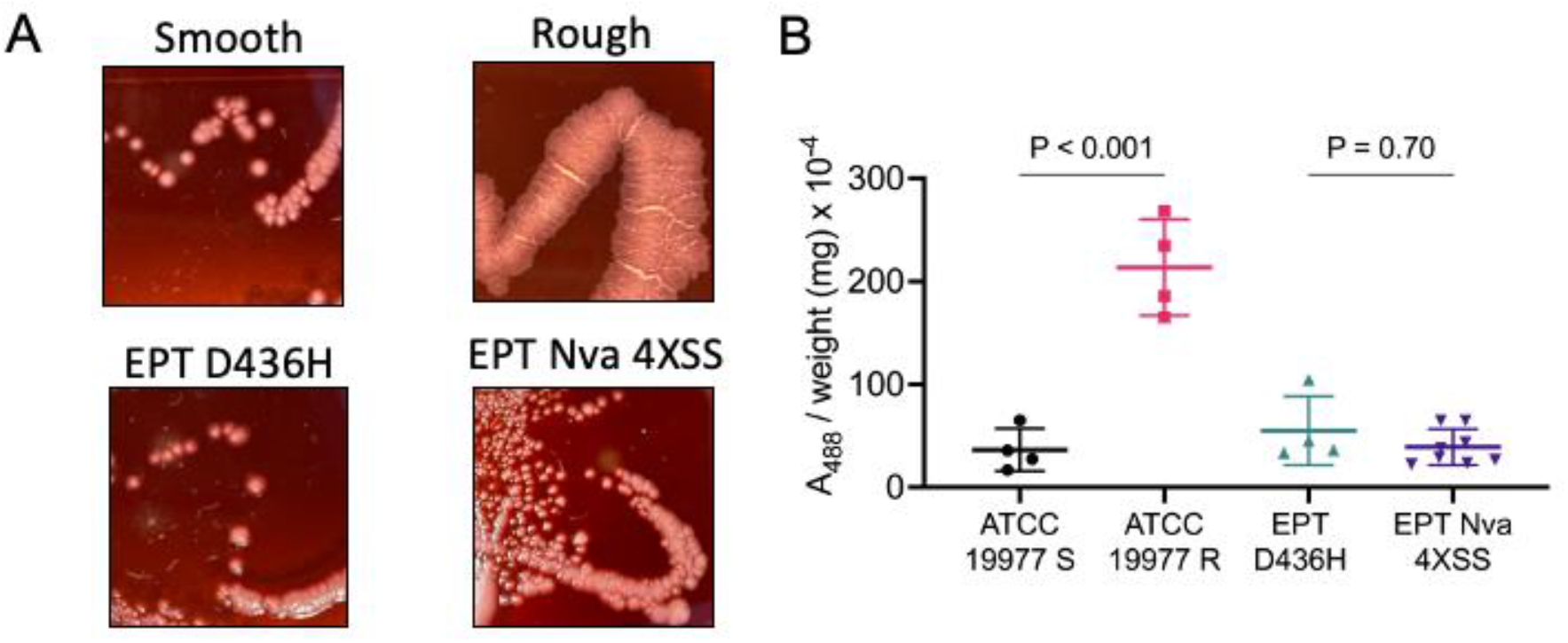
Membrane hydrophobicity of *M. abscessus*. (A) Colony morphology of ATCC 19977-S (smooth), ATCC 19977-R (rough), *M. abscessus* EPT^R^ D436H, and *M. abscessus* EPT^R^ Nva^R^ on 7H10 plates supplemented with 10% (v/v) OADC and Congo red at 140 μM. (B) Quantitative analysis of extracted Congo red dye retained in the membrane at A_488nm_. Data represents mean ± SD of n = 4 independent experiments with *M. abscessus* EPT^R^ Nva^R^ 4XSS in biological duplicates. P values were obtained by one-way ANOVA with Tukey’s multiple comparisons test.

### l-Norvaline Resistance is not Mediated by Efflux Pump Activity

Besides being involved in the architecture of the cell envelope some MmpL proteins have been shown to be antimycobacterial drug efflux pumps (34, 35). Most notably, this was demonstrated for the recently approved drug bedaquiline (36, 37). We speculated that the large number of efflux pumps in *M. abscessus* might contribute to l-norvaline resistance. To measure the net influx/efflux activity, we used an ethidium bromide (EtBr) accumulation assay previously used in *M. abscessus* and *M. tuberculosis* to assess drug resistance (38, 39). Initially, EtBr has limited fluorescence in aqueous solution but becomes appreciably fluorescent when intercalated with dsDNA (40). The fluorescence (Ex/Em 525/600) of intracellular EtBr was measured over 2 hours in wild type *M. abscessus* incubated with serial dilutions of EtBr (Fig 3*A*). Fluorescence time curves were fitted to an exponential plateau regression to determine the fluorescence at equilibrium (F_eq_) and generate a dose-response curve (Fig. 3*A* inset). The fluorescence of intracellular EtBr was measured in wild type, EPT^R^ LeuRS^D436H^, and EPT^R^ Nva^R^ 4XSS strains of *M. abscessus* at 6.25 μM EtBr. ATCC 19977 S and R morphotypes were used as controls. MmpL4b is non-functional in the rough morphotype, which resulted in impaired efflux and greater accumulation of EtBr (Fig. 3*B*). Although we measured a statistically significant difference in EtBr accumulation in the S morphotype compared to the R morphotype, the difference in F_eq_ between EPT^R^ Nva^R^ 4XSS and the parental strain was not statistically significant (Fig. 3*B*).

**Figure 3.**
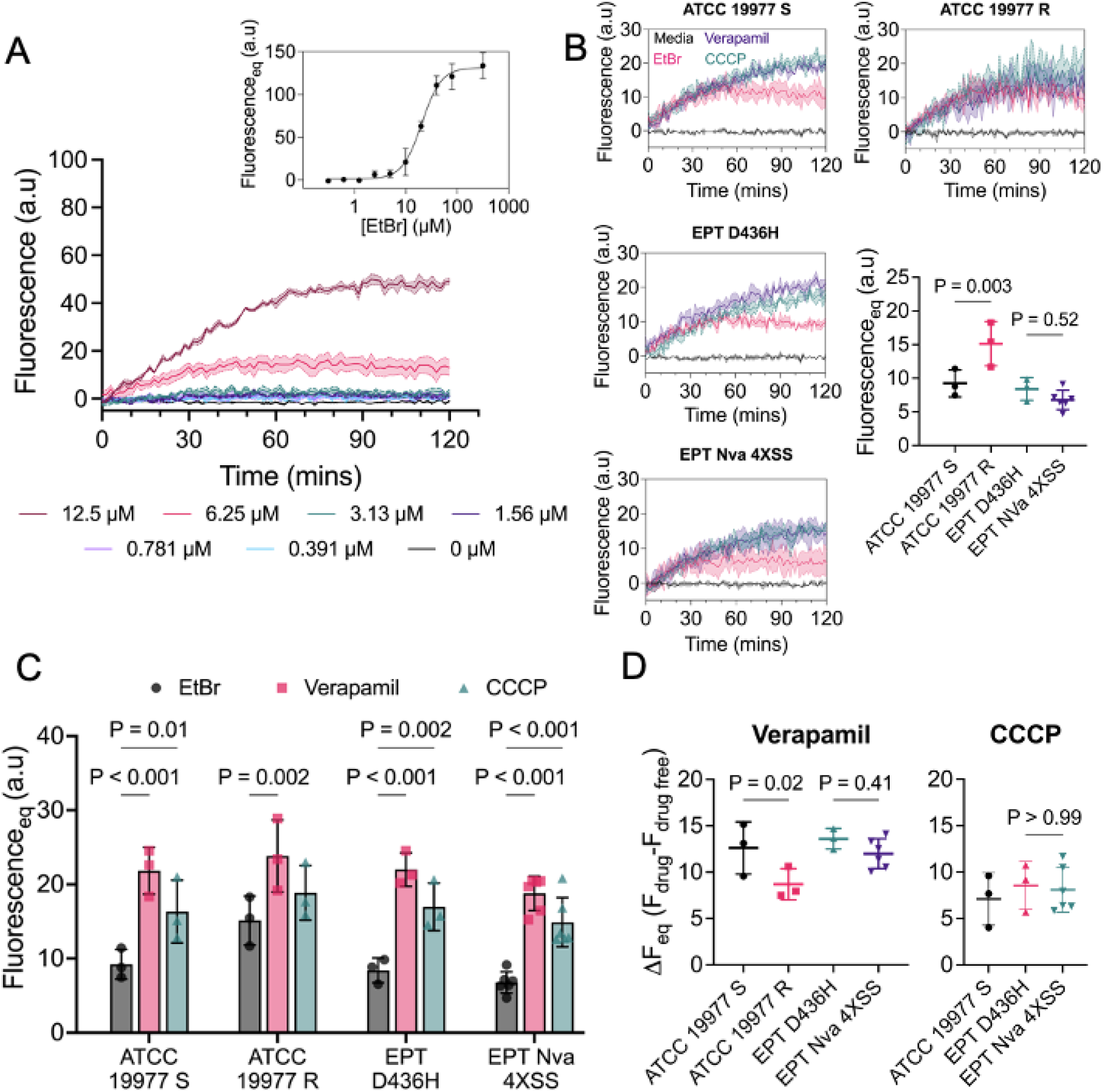
Efflux activity of *M. abscessus*. (A) Dose response curve of ethidium bromide (EtBr, Ex_525nm_, Em_600nm_) accumulation in *M. abscessus* ATCC 19977 S. (B) EtBr accumulation with 6.25 μM EtBr in ATCC 19977 S, ATCC 19977 R, *M. abscessus* EPT^R^ D436H, and *M. abscessus* EPT^R^ Nva^R^ 4XSS. 0.5 mM verapamil and 25 μM CCCP were used as efflux pump inhibitors in (B). Data is mean ± SD of n = 3 independent replicates with *M. abscessus* EPT^R^ Nva^R^ 4XSS in biological duplicates. P values were obtained by one-way ANOVA with Sidak’s multiple comparisons test. (C) Efflux pump inhibitors promote EtBr accumulation. Data is mean ± SD of n = 3 independent replicates with *M. abscessus* EPT^R^ Nva^R^ 4XSS in biological duplicates. P values were obtained by two-way ANOVA with Dunnett’s multiple comparisons test. (D) Quantification of efflux pump inhibitor effect. ATCC 19977 R was omitted from CCCP as the effect was not statistically significant. Data is mean ± SD of n = 3 independent replicates with *M. abscessus* EPT^R^ Nva^R^ 4XSS in biological duplicates. P values were obtained by one-way ANOVA with Sidak’s multiple comparisons test.

Next, we repeated the EtBr accumulation assay with verapamil and CCCP as efflux pump inhibitors (41). Using 0.5 mM verapamil and 25 μM CCCP, we measured a statistically significant increase in the F_eq_ for all strains except *M. abscessus* R with CCCP (Fig. 3*B* and *C*). With the efflux pump inhibitors validated, we sought to determine the effect size (ΔF_eq_ = F_drug_ – F_drug free_) of verapamil and CCCP on efflux pump inhibition. Verapamil and CCCP had similar effect sizes against EPT^R^ Nva^R^ 4XSS and the parental strain, which suggests that the EPT^R^ Nva^R^ 4XSS strain does not have increased efflux activity (Fig. 3*D*).

### α-IPMS^A555V^ Variant Participates in l-norvaline Resistance

To identify the putative mutation(s) underlying l-norvaline resistance, gDNA was extracted from *M. abscessus* EPT^R^ LeuRS^D436H^ as the parental strain and two *M. abscessus* EPT^R^ Nva^R^ 4XSS mutants for whole-genome sequencing. All strains sequenced retained the C to G transversion at position 1306 in *leuS* that translates into LeuRS^D436H^, which confirmed that resistance was not acquired through a D436H reversion as shown with the MIC to epetraborole (Fig. 1*A*). When compared to the parental strain, both EPT^R^ Nva^R^ 4XSS mutants sequenced had a unique C to T transition at position 1664 in *leuA* (MAB_0337c) and T to C transition at position 44 in tRNA^Leu(GAG)^ (MAB_t5031c). *leuA* codes for α-isopropylmalate synthase (α-IPMS) which catalyzes the carboxymethylation of 2-oxoisovalerate using acetyl-CoA to produce α-isopropylmalate as the first step in the l-leucine biosynthetic pathway. tRNA^Leu(GAG)^ is one of five encoded tRNA^Leu^ isoacceptors in *M. abscessus*.

To determine which variant gene is responsible for the l-norvaline resistant phenotype, wild type and mutant *leuA* and tRNA^Leu(GAG)^ were cloned with 250bp upstream to capture the nascent promoter into the integrative vector pMV306. These constructs, as well as an empty pMV306 vector (EV) were electroporated into *M. abscessus* EPT^R^ D436H. Next, we measured the bacterial viability of *M. abscessus* ATCC 19977, parental *M. abscessus* EPT^R^ D436H, complemented strains, and the natural double mutant *M. abscessus* EPT^R^ Nva^R^ 4XSS against l-norvaline, epetraborole, and amikacin. Only *M. abscessus* EPT^R^ D436H complemented with the mutant *leuA* had increased resistance to l-norvaline; complementation with the wild type *leuA* had no effect (Fig. S2*A*). All strains of *M. abscessus* with LeuRS^D436H^ retained resistance to epetraborole and all strains were equally susceptible to amikacin (Fig. S2*B* and *C*). l-norvaline resistance was not observed when *M. abscessus* EPT^R^ D436H was complemented with wild type or mutant *tRNA*^Leu(GAG)^ (Fig. S2*D-F*).

To corroborate the drug susceptibility results, we monitored the growth of *M. abscessus* ATCC 19977, *M. abscessus* EPT^R^ D436H, the *leuA* complemented strains and *M. abscessus* EPT^R^ Nva^R^ 4XSS in Sauton’s minimal media with 0.5 mM BCAAs. Complementing *M. abscessus* EPT^R^ D436H with the mutant variant of *leuA* nearly completely restored the growth in 0.5 mM l-norvaline relative to l-norvaline-free media while *M. abscessus* EPT^R^ D436H with wild type *leuA* or *M. abscessus* EPT^R^ D436H still exhibited reduced growth (Fig. 4*A* and Fig. S3). These results suggest that α-IPMS^A555V^ alone is sufficient to impart the l-norvaline resistant phenotype while the role of tRNA^Leu(GAG)T44C^ is yet to be determined.

**Figure 4.**
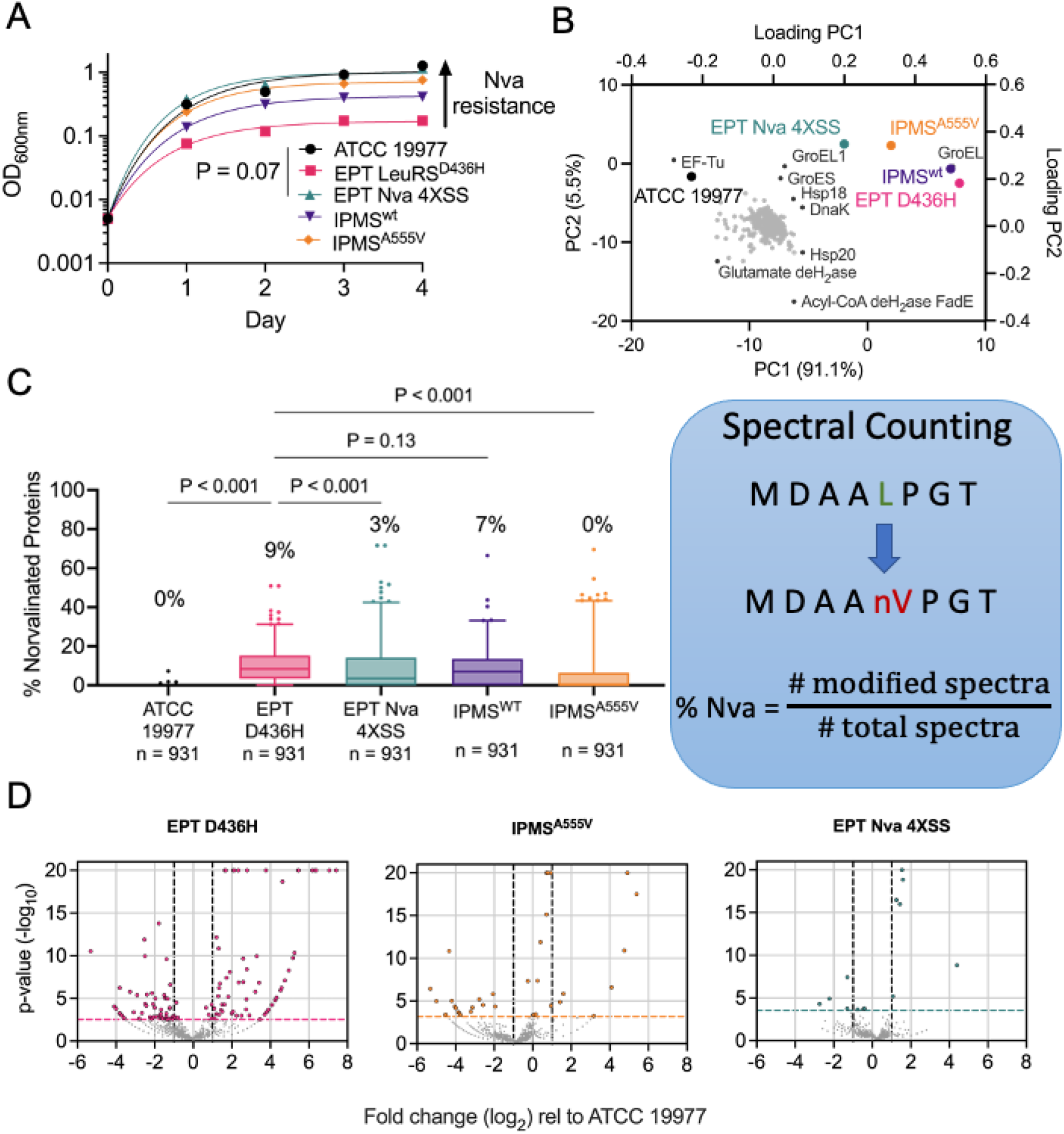
*M. abscessus* EPT^R^ Nva^R^ mutant uses alternative mechanism to limit l-norvaline toxicity. (A) Reference strain ATCC 19977 (black circles), EPT^R^ mutant LeuRS^D436H^ (pink squares), naturally raised EPT^R^ Nva^R^ mutant (green triangles), EPT^R^ mutant complemented with α-IPMS^WT^ (purple triangles) or α-IPMS^A555V^ (orange diamonds) grown in Sauton’s minimal media with 0.5 mM l-norvaline and fit with exponential plateau regression. Data is representative of n = 3 independent experiments with mean ± SD of technical triplicates. P values were obtained by one-way ANOVA with Tukey’s multiple comparisons test. (B) Principal component analysis of cell lysate proteomes after 24 h of growth in Sauton’s media with 0.5 mM l-norvaline. PC1 accounted for 90% of the variance, PC2 accounted for 5% of the variance. GroEL (MAB_0650) and Acyl-CoA dehydrogenase FadE (MAB_4437) had the greatest PC1 and PC2 coefficients, respectively. (C) Percentage of proteins with misincorporation of l-norvaline at leucine residues from cell lysates after 24 h of growth in Sauton’s media with 0.5 mM l-norvaline. Data is median with IQR, whiskers represent 1-99 percentile from spectral counting. P values were obtained by Friedman test with Dunn’s multiple comparisons test. (D) Volcano plots depict fold change (log_2_) and p-value(-log_10_) for each protein relative to the ATCC 19977 reference strain.

Previously, we showed that in the absence of LeuRS editing activity, l-norvaline is incorporated into proteins at sites coding for l-leucine (29). Therefore, we hypothesized that the *M. abscessus* EPT^R^ Nva^R^ 4XSS mutant would lack high levels of norvalinated proteins after l-norvaline challenge. Cell lysates were collected from *M. abscessus* ATCC 19977, *M. abscessus* EPT^R^ D436H, complemented strains, and *M. abscessus* EPT^R^ Nva^R^ 4XSS after 24 hours of growth in Sauton’s minimal media with 0.5 mM l-norvaline and analyzed by reverse-phase HPLC/MS. PCA of cell lysate proteomes identified three proteomic clusters among the five strains with PC1 and PC2 accounting for 91% and 5% of variance, respectively (Fig. 4*B* and S4*A*). Cluster 1 is the naturally l-norvaline resistant strain ATCC 19977, cluster 2 is the l-norvaline sensitive strain *M. abscessus* EPT^R^ D436H and its complement with wild type α-IPMS^WT^, and cluster 3 is the novel l-norvaline resistant strain *M. abscessus* EPT^R^ Nva^R^ 4XSS and the complement with α-IPMS^A555V^. The clusters were separated based on GroEL abundance in PC1 with a small contribution from Acyl-CoA dehydrogenase FadE abundance in PC2 (Fig. 4*B* and S4*B*). Proteins with l-norvaline residues were identified by filtering for l-leucine residues missing the 14 Da methylene group absent on l-norvaline. As expected from the PCA results, *M. abscessus* ATCC 19977 maintained preferential incorporation of l-leucine while the editing-deficient strain *M. abscessus* EPT^R^ D436H had a significant increase in norvaline misincorporation (Fig. 4*C* and S5*A*). *M. abscessus* EPT^R^ Nva^R^ 4XSS reduced the median level of norvaline misincorporation significantly despite lacking LeuRS editing activity (Fig. 4*C* and S5*A*). A decrease in median norvalination was also observed in the strain complemented with mutant *leuA* but not with the wild type *leuA* control (Fig. 4*C*). We also measured the level of l-norvaline in the culture supernatants after 24 hours of exposure and found no evidence of l-norvaline degradation or modification (Fig. S5*B*). To assess the overall bacterial stress response to l-norvaline challenge, we compared the proteomes of *M. abscessus* EPT^R^ D436H, complemented strains, and *M. abscessus* EPT^R^ Nva^R^ 4XSS to the ATCC 19977 reference strain. Higher levels of norvaline incorporation resulted in 134 statistically significant differentially expressed proteins (DEPs) (Fig. 4*D*). While *M. abscessus* EPT^R^ D436H had elevated levels of heat shock proteins and Clp proteases relative to the ATCC 19977 strain, we observed a significantly attenuated stress response in *M. abscessus* EPT^R^ Nva^R^ 4XSS with 10 DEPs and in *M. abscessus* EPT^R^ D436H complemented with α-IPMS^A555V^ with 28 DEPs (Fig. 4*D*).

### α-IPMS^A555V^ Mutation Located in Allosteric Site

To understand how the α-IPMS^A555V^ substitution resulted in minimal norvaline misincorporation across the proteome, we modelled α-IPMS_Mabs_ onto the experimentally determined structure of α-IPMS_Mtb_ using Swiss-Model (42). The current understanding is that the catalytic and regulatory domains are separated by a flexible hinge domain to accommodate conformational changes resulting from l-leucine bound in the allosteric site to elicit negative feedback inhibition (42). As a control, we compared the experimental structure of α-IPMS_Mtb_ (PDB 3FIG) to the α-IPMS_Mtb_ structure generated with Swiss-Model. We generated a homodimeric structure (QMEAN 0.87 ± 0.05) that was predicted to have the N-terminal catalytic (α/β)_8_ TIM barrel domain, a linker, and C-terminal regulatory domain with the βββα architecture (Fig. S6*A*). Next, we compared the overlaid structures of Swiss-Model generated α-IPMS_Mtb_ and α-IPMS_Mabs_ and observed substantial structural overlap with rmsd of 0.119 Å. From these results, we inferred homology between α-IPMS_Mtb_ and α-IPMS_Mabs_ and used the modelled α-IPMS_Mabs_ structure to hypothesize the effect of α-IPMS^A555V^ substitution identified in *M. abscessus* EPT^R^ Nva^R^ 4XSS.

The experimentally determined structure of α-IPMS_Mtb_ was solved as a homodimer with an N-terminal catalytic domain and a C-terminal regulatory domain (PDB 3FIG and 3HPZ). The regulatory domains contain two symmetrical pockets at the dimer interface with loops in an open conformation when unbound to l-leucine (PDB 3HPZ and Fig. 5*B*) (42). However, the loop adopts a closed conformation when l-leucine is bound in the hydrophobic pocket of the regulatory domain (PDB 3FIG, Fig. 5*C*) (42). The A555V substitution identified from WGS in *M. abscessus* EPT^R^ Nva^R^ 4XSS is located on this loop (Fig. 5*C*). We modelled the A555V substitution on α-IPMS_Mabs_ and observed a deviation > 4*σ* from the ideal angle with D557 and S554 was identified as a rotamer outlier. Taken together with the additional electron density from A555V next to the side chain of L546, we hypothesized that A555V would prevent the loop from adopting a closed conformation when bound by l-leucine and disrupt the allosteric feedback inhibition.

**Figure 5.**
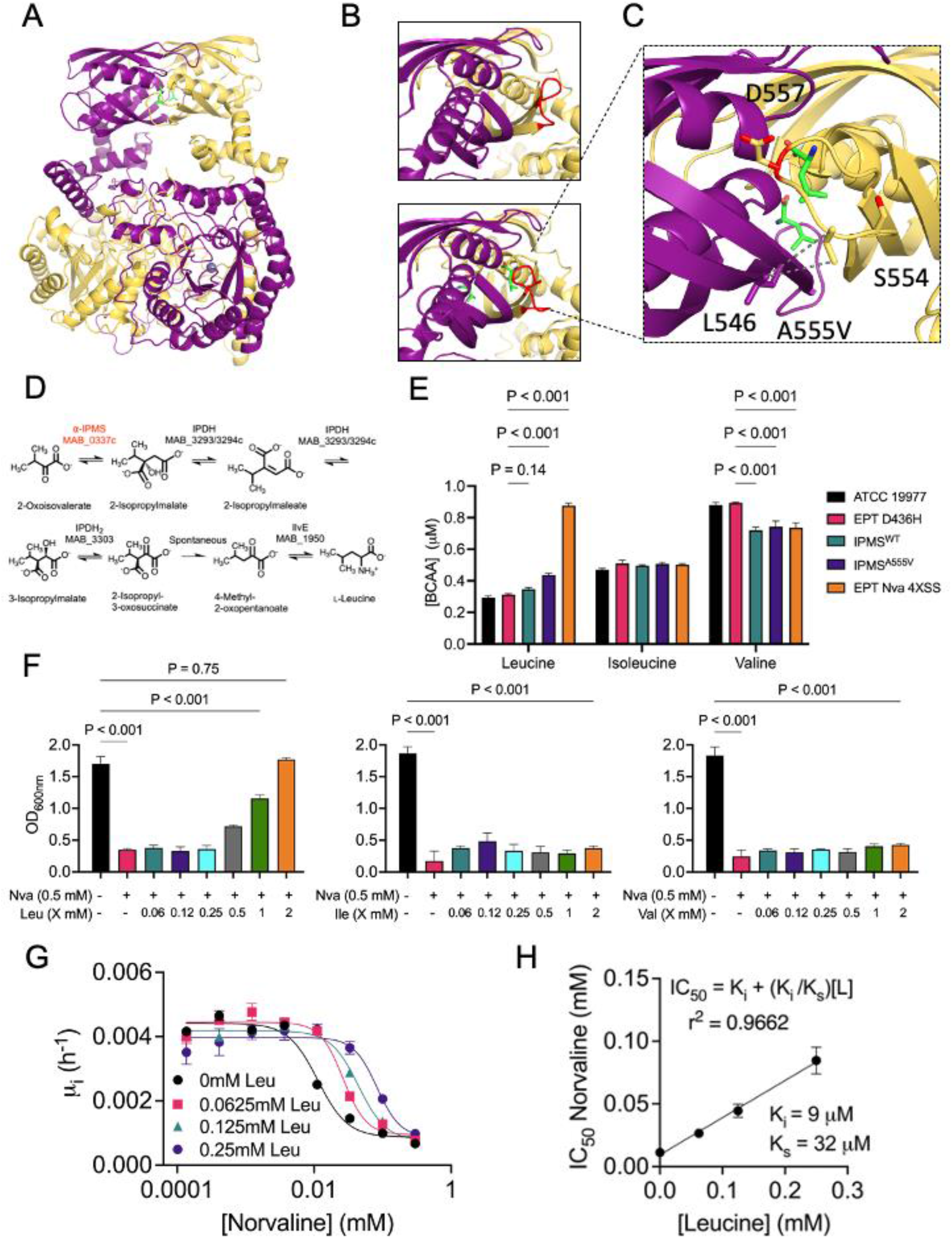
α-IPMS^A555V^ variant over produces l-leucine to outcompete l-norvaline. (A) Swiss-Model structure of α-IPMS_Mabs_ from *M. tuberculosis* PDB 3FIG (42). Monomers in yellow and purple, l-leucine in green, and Zn^2+^ as blue spheres. (B, top) C-terminal regulatory domain without l-leucine bound in an open conformation (red loop). (B, bottom) C-terminal regulatory domain with l-leucine bound in a closed conformation. (C) Effects of mutated A555V highlighted. (D) Pathway for l-leucine synthesis. (E) BCAA concentrations in culture filtrates of reference strain ATCC 19977, EPT^R^ mutant LeuRS^D436H^, EPT^R^ mutant complemented with IPMS^WT^ or IPMS^A555V^, and naturally raised EPT^R^ Nva^R^ mutant grown in Sauton’s minimal media. Data is mean ± SD of n = 3 biological replicates. P values were obtained by one-way ANOVA with Tukey’s multiple comparisons test. (F) *M. abscessus* EPT^R^ D436H strain grown in Sauton’s minimal media ± l-norvaline and serial dilutions of l-leucine, l-isoleucine, or l-valine. Data is mean ± SD of n = 3 independent experiments. P values were obtained by one-way ANOVA with Tukey’s multiple comparisons test. (G) Initial growth rates (first 24 h) of *M. abscessus* EPT^R^ D436H in l-norvaline supplemented with l-leucine. (H) l-norvaline inhibition fit to a model of competitive inhibition. K_i_ is the inhibition constant of l-norvaline. K_s_ is the substrate constant for l-leucine. L is the concentration of l-leucine.

### Overproduction of l-Leucine Competitively Inhibits Norvaline

To understand the mechanism of l-norvaline resistance in the absence of LeuRS editing activity, we hypothesized that the mutation in α-IPMS would disrupt the equilibrium of the l-leucine biosynthetic pathway and confer abnormal l-leucine production through loss of allosteric regulation in α-IPMS (Fig. 5*D*). To test this hypothesis, we grew *M. abscessus* ATCC 19977, *M. abscessus* EPT^R^ D436H, complemented strains, and *M. abscessus* EPT^R^ Nva^R^ 4XSS in Sauton’s minimal media and measured the concentration of BCAAs in the culture filtrate using HPLC/MS. Indeed, *M. abscessus* strains that harbored α-IPMS^A555V^ produced higher concentrations of l-leucine relative to the ATCC 19977 reference strain while l-isoleucine concentrations were unaffected (Fig. 5*E*). l-Valine concentrations, however, were significantly reduced in strains that produced excess l-leucine (Fig. 5*E*).

Next, we examined the rescue effect of l-leucine by growing *M. abscessus* EPT^R^ D436H in 0.5 mM l-norvaline with increasing l-leucine concentrations. After day 7 of growth, we observed that a 1:1 ratio of l-leucine:l-norvaine restored the growth of *M. abscessus* EPT^R^ D436H to 50% of the l-norvaline free conditions while l-isoleucine and l-valine had no rescue effect (Fig. 5*F*). Growth was completely restored with a 4:1 ratio of l-leucine:l-norvaline (Fig. 5*F*). Motivated by these results, we asked whether l-leucine could act as a competitive inhibitor to l-norvaline. We measured the initial growth rates from time 0 to 24 hours in serial dilutions of l-norvaline to generate a dose-response curve (Fig. 5*G*). These measurements were repeated with increasing l-leucine concentrations and required higher concentrations of norvaline to generate saturating growth inhibition. To obtain IC_50_ values for l-norvaline, the saturating curves were fit to four parameter dose-response regression. Next, IC_50_ values were used to estimate an apparent inhibition constant of l-norvaline, K_i_, and an apparent substrate constant of l-leucine, K_s_, using the Cheng-Prusoff theorem for competitive inhibition (Fig. 5*H*). The linear relationship between the IC_50_ of l-norvaline and the concentration of l-leucine suggests competitive inhibition between the amino acids where l-norvaline (K_i_ = 9 μM) is 3-4x more potent as an inhibitor than l-leucine (K_s_ = 32 μM) is as a substrate (Fig. 5*H*).

## Discussion

*M. abscessus* infections are notoriously difficult to treat largely due to its enhanced toolkit of antimicrobial resistance mechanisms (6). We show that *M. abscessus* can subvert the toxicity of l-norvaline, a non-canonical amino acid shown to be an experimental adjunct agent with epetraborole (29). Classically, *M. abscessus* is naturally resistant to l-norvaline because of the editing domain on LeuRS. LeuRS provides pre-transfer editing before the incorrect amino acid is transferred to tRNA^Leu^ or post-transfer editing when the erroneous amino acid has already been transferred onto tRNA^Leu^ (43). In both mechanisms, the editing activity is mediated by a conserved aspartic acid residue, D436. Our results demonstrate that a *M. abscessus* mutant can limit the incorrect use of l-norvaline in the absence of pre- and post-transfer editing activity. We show that resistance is not the result of modified membrane hydrophobicity nor increased efflux activity but stems from the loss of allosteric regulation to overproduce a metabolite and outcompete the inhibitor.

The A555V mutation was found in α-IPMS, the first enzyme involved in l-leucine biosynthesis (44). α-IPMS was previously shown to have a regulatory domain with a hydrophobic pocket where l-leucine binds with the carboxylate positioned between the partial positively charged N-terminal ends of two helices, similar to the bound chloride ion in the absence of l-leucine (42). From the crystal structure, it was hypothesized to be a regulatory domain for feedback inhibition. Later, kinetic evidence emerged that supported intradomain communication by an allosteric mechanism with l-leucine (45). This regulation had also been postulated in yeast where mutations were identified in the C-terminal regulatory domain that resulted in insensitivity to l-leucine feedback inhibition (46). Other non-regulatory yeast α-IPMS mutants became resistant to Zn^2+^-mediated inactivation by Coenzyme-A. This was later supported with kinetic data examining the effects of divalent cations on the activity of α-IPMS as a mechanism to control energy metabolism (46, 47).

In line with the allosteric mechanism of α-IPMS, our results show an increased production of l-leucine by strains harboring α-IPMS^A555V^ relative to the ATCC 19977 reference strain. We also tested l-isoleucine and observed no difference in BCAA production. l-Valine, however, was significantly reduced in strains that produced more l-leucine. Although BCAAs share structural features, the biosynthetic pathway for l-isoleucine is dependent on l-threonine production while l-leucine/l-valine are synthesized from pyruvate. The α-IPMS negative feedback mechanism is believed to be specific for l-leucine since α-IPMS catalyzes the first committed step of l-leucine biosynthesis after diverging away from the shared l-valine pathway (44). l-Isoleucine and l-valine have also been shown to be allosteric inhibitors and activators, respectively, but towards the l-isoleucine chokepoint enzyme, threonine deaminase (48). Unregulated α-IPMS activity that shifts the equilibrium of 2-oxoisovalerate could explain why l-valine production was stunted in strains with α-IPMS^A555V^.

*M. abscessus* grown under l-norvaline challenge showed that α-IPMS^A555V^ strains were more resistant to growth inhibition than α-IPMS^WT^ strains. Strains with LeuRS^D436H^ fail to grow in 0.5 mM l-norvaline however, the natural double mutant *M. abscessus* EPT^R^ Nva^R^ 4XSS and *M. abscessus* EPT^R^ D436H complemented with α-IPMS^A555V^ had comparable growth to *M. abscessus* ATCC 19977. These data suggest that overproduction of l-leucine from the mutation in the loop capping the allosteric site results in resistance to l-norvaline. The growth kinetics of *M. abscessus* EPT^R^ D436H in l-norvaline and l-leucine shed light on the requirement for additional l-leucine and the biochemical basis for l-leucine rescue. By measuring initial growth rates, we observed increasing l-norvaline IC_50_ values proportional to the l-leucine concentration, suggestive of competitive inhibition with K_i_ = 9 μM for l-norvaline and K_s_ = 32 μM for l-leucine. The inhibition and substrate constants lend additional support to the l-leucine rescue experiments where nearly 4x more l-leucine was required to fully restore the growth of *M. abscessus* EPT^R^ D436H in 0.5 mM l-norvaline.

Corroborating our previous results, strains that produced higher levels of l-leucine and misincorporated less l-norvaline in the proteome had superior growth output in l-norvaline. The proteomic profiles suggest that strains with α-IPMS^A555V^ can tolerate l-norvaline as the unfolded protein response observed in *M. abscessus* EPT^R^ D436H, characterized by heat shock proteins and chaperonins in the dominant principal component 1, is diminished in *M. abscessus* EPT^R^ Nva^R^ 4XSS. The variance in principal component 2 is attributed to an increased FadE abundance in strains that failed to grow during norvaline challenge. FadE is predicted to be an acyl-CoA dehydrogenase involved in fatty acid β-oxidation, which suggests that *M. abscessus* EPT^R^ D436H and α-IPMS^wt^ complemented strain upregulated fatty acid β-oxidation to meet the metabolic requirements of misfolded protein proteolysis. Overall, the proteome of strains with α-IPMS^A555V^ clustered separately from *M. abscessus* EPT^R^ D436H and appeared more similar to the ATCC 19977 reference strain.

In conclusion, we describe a novel mechanism of resistance to an antimicrobial agent. Systematic evaluation revealed that the *M. abscessus* resistance phenotype was due to the loss of allosteric feedback inhibition in a biosynthetic enzyme. *M. abscessus* expressing α-IPMS^A555V^ gained the ability to produce excess quantities of l-leucine, with no deleterious consequences, to outcompete l-norvaline from being used as a LeuRS substrate. This mutation reduces norvalination of the proteome and toxicity associated with misfolded proteins. Our work here provides evidence for a mechanism of antimicrobial resistance in *M. abscessus* where an orthogonal mutation in an allosteric chokepoint enzyme rescues the bacteria through dysregulation of BCAA metabolism.

## Materials and Methods

### Culture conditions

*M. abscessus* strains were grown in rolling liquid culture at 37 °C in Middlebrook 7H9 (Difco) supplemented with 10% (v/v) albumin dextrose catalase (ADC), 0.2% (v/v) glycerol, and 0.05% (v/v) Tween 80 (7H9 complete) or Sauton’s defined minimal media. 7H10 agar plates supplemented with 10% (v/v) oleic acid ADC (OADC) and 0.5% (v/v) glycerol were used for growth on solid media at 37 °C.

### Norvaline time-dependent killing

Log phase (OD_600_ of 0.4-0.8) *M. abscessus* was diluted to an OD_600_ of 0.0001 (10^5^ CFU/mL) and challenged with an initial dose of l-norvaline at 4X MIC_90_ of the sensitive *M. abscessus* LeuRS^D436H^ strain or norvaline at 4X MIC_90_ supplemented daily to create steady-state conditions (4XSS). One hundred μL aliquots were taken and serially diluted in 7H9 complete and plated on 7H10 agar. The starting inoculum was determined from time 0 before norvaline was added. CFUs were determined after 4 days of incubation at 37 °C. Bactericidal activity is defined as a 3 log_10_ decrease (99.9%) from the starting inoculum.

### Minimum inhibitory concentrations (MIC)

MIC values were determined using the resazurin microtiter assay (REMA). Cultures were grown to log phase (OD_600_ of 0.4-0.8) and diluted to OD_600_ of 0.005 (5×10^6^ CFU/mL). Compounds were prepared in two-fold serial dilutions in 96-well plates with 90 μL of bacteria per well to a final volume of 100 μL. 96-well clear, flat bottom plates were incubated at 37 °C until drug-free wells were turbid (2 days for *M. abscessus*). Ten μL of resazurin prepared at 0.025% (w/v) in distilled water was added to each well. Once the drug-free wells turned pink (3-4 hours or one doubling time), the fluorescence (Ex/Em 560/590) was measured using an Infinite F200 Tecan plate reader. Fluorescence intensities were converted to percentage of viable cells relative to drug-free conditions and fit to the modified four-parameter dose-response regression using GraphPad Prism version 9. MIC values at 90% reduction of viability were determined from the Gompertz nonlinear regression equation (49).

### Isolating l-norvaline resistant mutants

l-norvaline resistant mutants of *M. abscessus* ATCC 19977 were isolated from l-norvaline time-dependent killing assays. Single colonies were obtained from streaked 7H10 plates. Genomic DNA (gDNA) was extracted from *M. abscessus* EPT^R^ LeuRS^D436H^ as the parental strain and two colonies of *M. abscessus* EPT^R^ Nva^R^ 4XSS mutants with confirmed norvaline resistance (REMA method) using the Qiagen QiaAMP UCP Pathogen mini kit with a modified mechanical lysis protocol. Culture was pelleted by centrifugation and resuspend in 590 μL of ATL buffer containing the Dx reagent in a low-bind tube. 40 μL of proteinase K (20 mg/mL) and 20 μL of lysozyme (100 mg/mL) were added and incubated for 30 minutes at 56 °C under agitation. Solution was transferred into a Pathogen Lysis Tube L and vortexed twice using a FastPrep-24 instrument at 6.5 m/s for 45 s with a 5-minute incubation on ice in between. Supernatant was transferred into a fresh 2 mL low-bind tube and gDNA was collected via spin protocol according to manufacturer’s instructions. gDNA was quantified with Quant-iT PicoGreen dsDNA kit.

### Sequencing and variant calling

Paired-end read sequencing libraries were prepared using Illumina S4 kit. Whole genome sequencing was performed on Illumina Novaseq 6000. Raw paired-end reads were adapter and quality trimmed with Trimmomatic v0.40 and aligned to the *M. abscessus* ATCC 19977 reference genome (GenBank: CU_458896.1) using Burrows-Wheeler Aligner - maximum exact matches algorithm (bwa-mem) (50, 51). Aligned reads were sorted using SAMTOOLs (52). Duplicate reads were filtered using Picard. Freebayes v1.3.6 was used for variant calling and filtering variants with mapping quality 60, minimum coverage 10, minimum allele frequency of 0.5, and QUAL >100 (53). Variants identified in parental and resistant strains were manually compared to isolate variants unique to resistant strains.

### Growth curves

Bacteria were grown to log phase (OD_600_ of 0.4-0.8) in 7H9 complete or Sauton media and diluted to OD_600_ of 0.005 (5×10^6^ CFU/mL). Branched-chain amino acids were added to 90 μL of bacteria to a final volume of 100 μL. Growth curves were performed in 96-well clear, flat bottom microtiter plates, incubated statically at 37 °C before measuring A_600_ on an Infinite F200 Tecan plate reader. Data was fit to an exponential plateau regression to determine the growth rate μ and maximum growth plateau Y_M_.

### Congo red uptake

Strains were grown on 7H10 agar plates supplemented with 10% (v/v) OADC, 0.5% (v/v) glycerol, and Congo red at 140 μM at 37 °C. After five days, cells were scraped into an Eppendorf tube and washed with water until the supernatant turned clear. Washed cells were incubated with 1 mL of DMSO for 2 hours. Supernatants were collected and measured at A_488_ using an Infinite F200 Tecan plate reader. Values were normalized to the dry weight of the pellet.

### EtBr efflux

*M. abscessus* strains were grown to log phase (OD_600_ of 0.4-0.8) in Sauton media and diluted to OD_600_ of 0.2. 80 μL of bacteria was added to a black, clear bottom 96-well plate with 10 μL serially diluted ethidium bromide and 10μL of media. For efflux pump inhibition assays, media was replaced with 10 μL of verapamil or CCCP for a final concentration of 0.5 mM or 25 μM, respectively. Fluorescence (Ex/Em 525/600) was measured every 90 seconds for 2 hours at 25 °C using an Infinite F200 Tecan plate reader.

### Cloning and over-expressing *leuA* in *M. abscessus* EPT^R^ LeuRS^D436H^

The wildtype and mutant *leuA* gene (MAB_0337c), which encodes α-isopropylmalate synthase, and tRNA^Leu(GAG)^ gene (MAB_t5031c) including 250bp upstream to capture the native promoter were PCR amplified from gDNA using Q5 polymerase with forward primer (5’ ACTGCAGAATTCTCCTTGAGAGCGCTCGCGAAG) and reverse primer (5’ GATGATAAGCTTCTAGGCGCGAGCAGCACGG), ligated into pMV306_hsp60 digested with EcoRV and HindIII restriction enzymes, and transformed into *E. coli* DH5*α* cells (Promega). Plasmids were extracted and sequenced using Sanger sequencing.

### Initial growth kinetics and data theory

*M. abscessus* strains were grown to log phase (OD_600_ of 0.4-0.8) in Sauton media and diluted to OD_600_ of 0.005 in 96-well clear, flat bottom microtiter plates with serial dilutions of norvaline. Each set of norvaline dose-response curves was incubated with a constant amount of l-leucine as a norvaline inhibitor. Plates were incubated statically at 37 °C and A_600_ was measured on an Infinite F200 Tecan plate reader to monitor growth. Initial growth rates, μ_i_ (h^-1^), were measured during the first 24 hours of growth. Growth rate inhibition, performed under variable norvaline concentrations and constant l-leucine concentrations (0 mM, 0.0625 mM, 0.125 mM, or 0.250 mM) were fitted to a dose-response curve

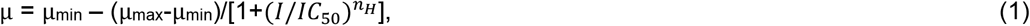

where μ is the rate in the presence of inhibitor at concentration *I*, μ_max_ and μ_min_ are the maximum and minimum growth rates, *IC*_50_ is the concentration of inhibitor that inhibits the growth rate by 50%, and *n*_H_ is the Hill coefficient. *IC*_50_ values were used to estimate an apparent inhibition constant, *K*_i_, using the Cheng-Prusoff equation for competitive inhibition

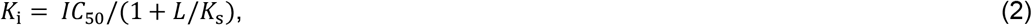

where *K*_i_ is the apparent inhibition constant for norvaline, *L* is the concentration of l-leucine, and *K*_s_ is the substrate affinity constant of l-leucine.

### LC-MS/MS measurement of norvaline in the proteome

*M. abscessus* strains were grown to early log phase (OD_600_ of 0.1-0.2) and challenged with 0.5 mM l-norvaline in Sauton media for 24 hours. Proteins were extracted using an optimized protocol for mass spectrometry follow-up (54). Cells were collected, washed with ice-cold PBS, and resuspended in 1mL lysis buffer (50 mM NH_4_HCO_3_ pH 7.4, 10 mM MgCl_2_, 0.1% NaN_3_, 1 mM EGTA, 1 x protease inhibitors (Roche), 7 M urea, and 2 M thiourea). Cells were lysed with zirconia beads and the cell lysate was collected and filtered through a 0.22 μm membrane. Proteins were precipitated overnight at 4 ºC with TCA at 25% (v/v). The precipitate was washed with 1 mL cold acetone and 250 μL cold water. The final wash is only water. The pellet was resuspended in 200 μL resuspension buffer (50 mM NH_4_HCO_3_, 1 M urea). Protein extraction was quantified with the Bradford assay and the quality of proteins was examined on SDS-PAGE. Protein lysates were dissolved in SDS-PAGE reducing buffer and electrophoresed onto a single stacking gel band to remove lipids, detergents and salts. For each sample, a single gel band was reduced with DTT (Sigma), alkylated with iodoacetic acid (Sigma) and digested with LCMS grade trypsin (Promega). Extracted peptides were re-solubilized in 0.1% aqueous formic acid and loaded onto a Thermo Acclaim Pepmap (Thermo, 75 μM ID X 2 cm C18 3 μM beads) precolumn and then onto an Acclaim Pepmap Easyspray (Thermo, 75 μM X 15 cm with 2 μM C18 beads) analytical column separation using a Dionex Ultimate 3000 uHPLC at 250 nl/min with a gradient of 2-35% organic (0.1% formic acid in acetonitrile) over 2 hours. Peptides were analyzed using a Thermo Orbitrap Fusion mass spectrometer operating at 120,000 resolution for MS1 with HCD sequencing at top speed (15,000 resolution) for all peptides with a charge of 2+ or greater. The raw data were converted into *.mgf format (Mascot generic format) for searching using the Mascot 2.5.1 search engine (Matrix Science) against *M. abscessus* protein sequences (Uniprot downloaded 2020.11.30). A modification for Xle->Val was used to detect incorporation of Val into wild-type sequences. The database search results were loaded onto Scaffold Q+ Scaffold_4.4.8 (Proteome Sciences) for statistical treatment and data visualization.

### Principal component analysis

The dataset comprised of the 1000 most abundant proteins from each strain of *M. abscessus* based on quantitative spectral counts. Spectral counts for each protein were standardized within strains. PCA was performed using R to generate the loading data and scores. PCA was visualized with the first two principal components which accounted for >95% of the variance.

### LC-MS measurement of BCAA

Samples aliquots were diluted with an internal standard (norvaline-d5) and derivatized with 6-aminoquinolyl-N-hydroxysuccinimidyl carbamate (AQC; Toronto Research Chemicals, Ontario, Canada) for analysis using reversed phase ultra performance liquid chromatography mass spectrometry (UPLC-MS). Samples were prepared and analyzed along with calibration curves containing isoleucine, leucine, norvaline and valine (Sigma-Aldrich, St. Louis, Missouri, USA) in culture media. An internal standard working solution (ISWS) containing 50 μM norvaline-d5 (CDN Isotopes, Dorval, Quebec, Canada) in water was added to the experimental and calibration samples prior to derivatization. ISWS aliquots (25 μL) were added to sample aliquots (25 μL) in microcentrifuge tubes and vortexed. Aliquots (10 μL) then were transferred into glass tubes containing 70 μL buffer solution (0.2M sodium borate pH 8.8) along with 20 μL derivatization solution (10mM AQC in acetonitrile), mixed and incubated for 10 min at 55°C. After cooling to room temperature, aliquots (5 μL) were transferred to autosampler vials containing 995 μL Type-1 water for UPLC-MS analysis. Samples were analyzed by UPLC-MS using an Agilent 6460 triple quadrupole mass spectrometer coupled with an Agilent 1290 UPLC system (Agilent, Santa Clara, California, USA). Samples (5μL) were injected onto an Agilent Eclipse Plus C18 100 × 2.1 mm (1.8 μm) column and chromatographed with a reverse phase gradient at 0.250 mL/min using 0.1% formic acid in water and 0.1% formic acid in acetonitrile. The derivatized amino acids were detected using electrospray positive mode ionization followed by MS/MS fragmentation. Data acquisition was performed using Agilent MassHunter Data Acquisition (version B.04.01) software. Peak area measurements from selected product ions, calibration curve regression analysis and resulting sample quantification were performed using Agilent MassHunter Quantitative Analysis (version B.05.00) software.

### Homology modeling

The models of α-IPMS_Mabs_ (MAB_0337c) were generated using the SWISS-MODEL server and the crystal structures of α-IPMS_Mtb_ as templates (PDB 3FIG, 3HPZ) (55, 56).

### Data analysis

Data were processed by GraphPad PRISM 9.4.1, The PyMOL Molecular Graphics System 2.3.0, R 4.1.2, and Scaffold Q+ Scaffold_4.4.8.

## Acknowledgments

We thank the entire team at the Clinical Proteomics Platform of the Research Institute of the McGill University Health Centre [RI-MUHC] for discussion and their assistance with mass spectrometry. Work in M.A.B.’s laboratory was supported by a Foundation Grant from the Canadian Institutes for Health Research (FDN-148362). J.R.S. was supported by a doctoral studentship from the Fonds de recherche du Québec. M.A.B. is the recipient of a Tier 1 Canada Research Chair.

## Supporting Information

**Fig. S1.**
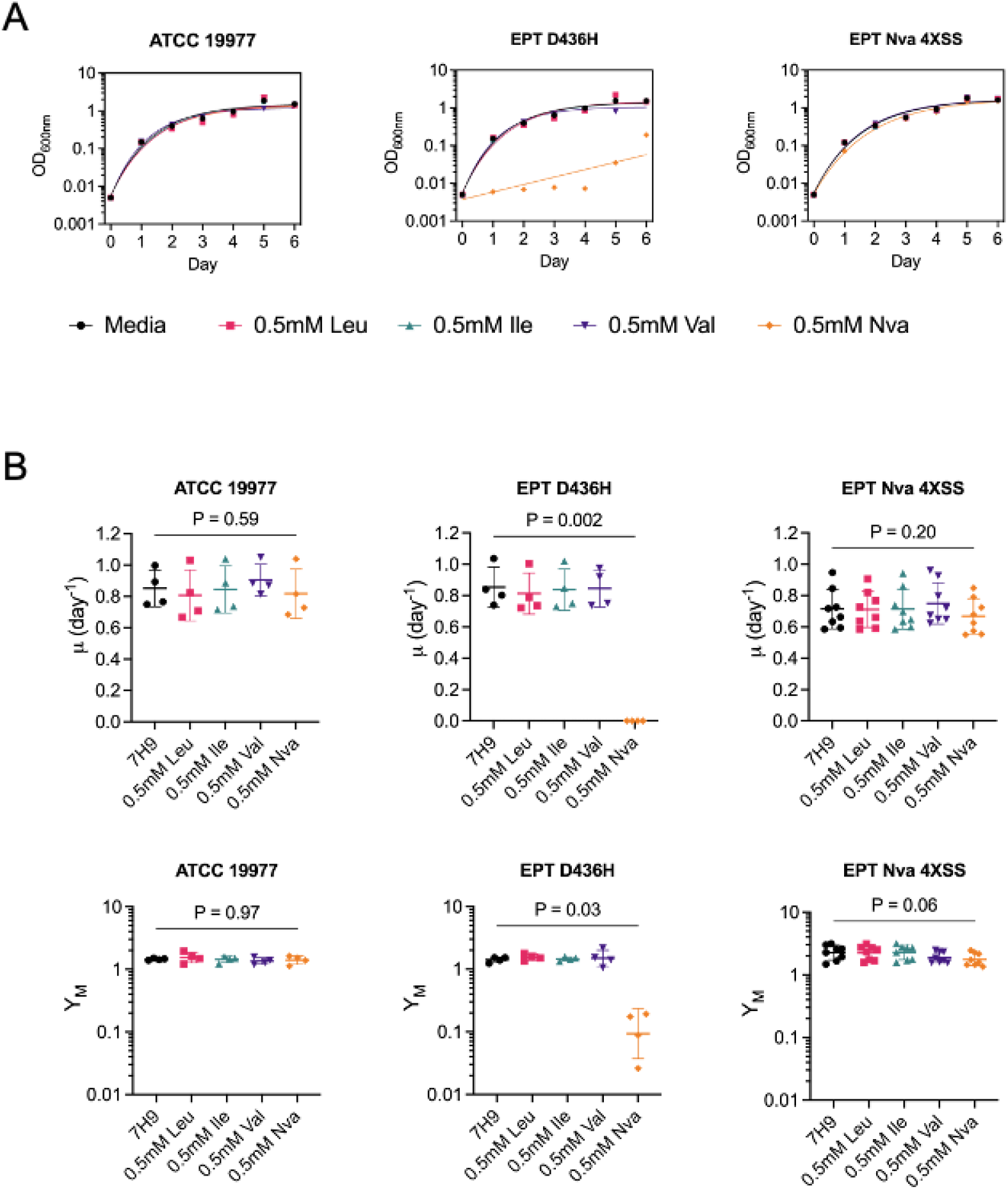
Growth characteristics of *M. abscessus* EPT^R^ Nva^R^ mutant in 7H9 media. (A) Growth curves of *M. abscessus* ATCC 19977, EPT^R^ *M. abscessus* with LeuRS^D436H^, and a mutant with dual EPT^R^ and Nva^R^ resistance raised at 4X MIC_90_ SS. Strains were grown statically in 96-well plates with nutrient rich 7H9 media ± BCAAs and fit with exponential plateau regression. Data is representative of n = 4 independent experiments with mean ± SD of technical triplicates. (B) Comparison of growth rates (**μ**) and maximum growth plateau (Y_M_) in 7H9 media ± BCAAs. Data is mean ± SD from n = 4-8 independent replicates. P values were obtained by one-way ANOVA with Dunnett’s multiple comparisons test.

**Fig. S2.**
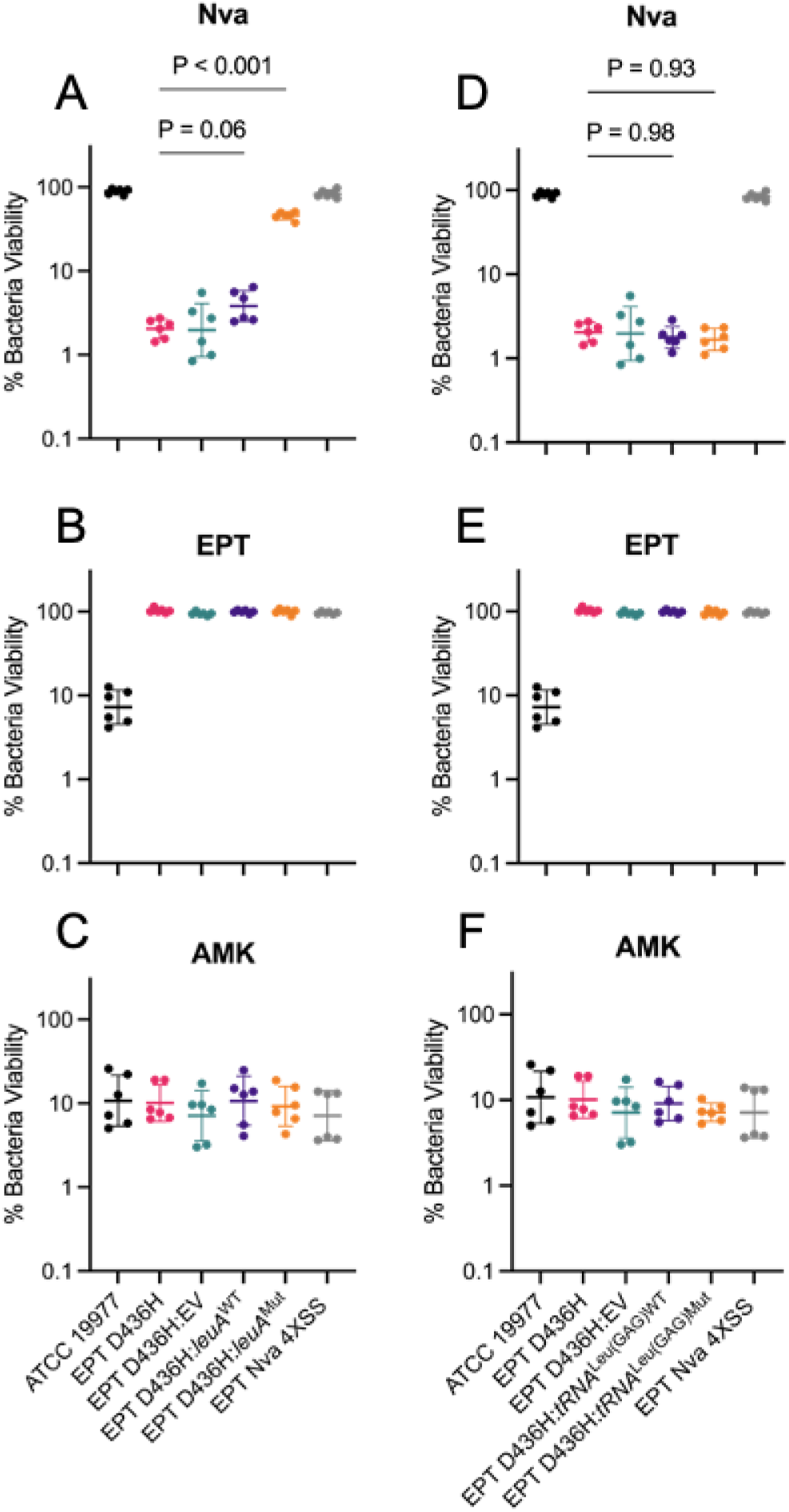
*M. abscessus* EPT^R^ D436H strain complemented with putative *leuA* or *tRNA*^Leu(GAG)^ resistance variants. Reference strain ATCC 19977 (black), EPT^R^ mutant LeuRS^D436H^ (pink), EPT^R^ mutant complemented with empty pMV306 vector (green), EPT^R^ mutant complemented with wild type *leuA* (purple) or complemented with mutant *leuA* (orange), and naturally raised EPT^R^ Nva^R^ mutant (grey) grown in (A) l-norvaline, (B) epetraborole, or (C) amikacin. Alternatively, the EPT^R^ LeuRS^D436H^ mutant was complemented with wild type *tRNA*^leu^ (purple) or mutant *tRNA*^leu^ (orange) and grown in (D) l-norvaline, (E) epetraborole, (F) amikacin. Data is n = 2 independent experiments with mean ± SD of technical triplicates. P values were obtained by one-way ANOVA with Tukey’s multiple comparisons test.

**Fig. S3.**
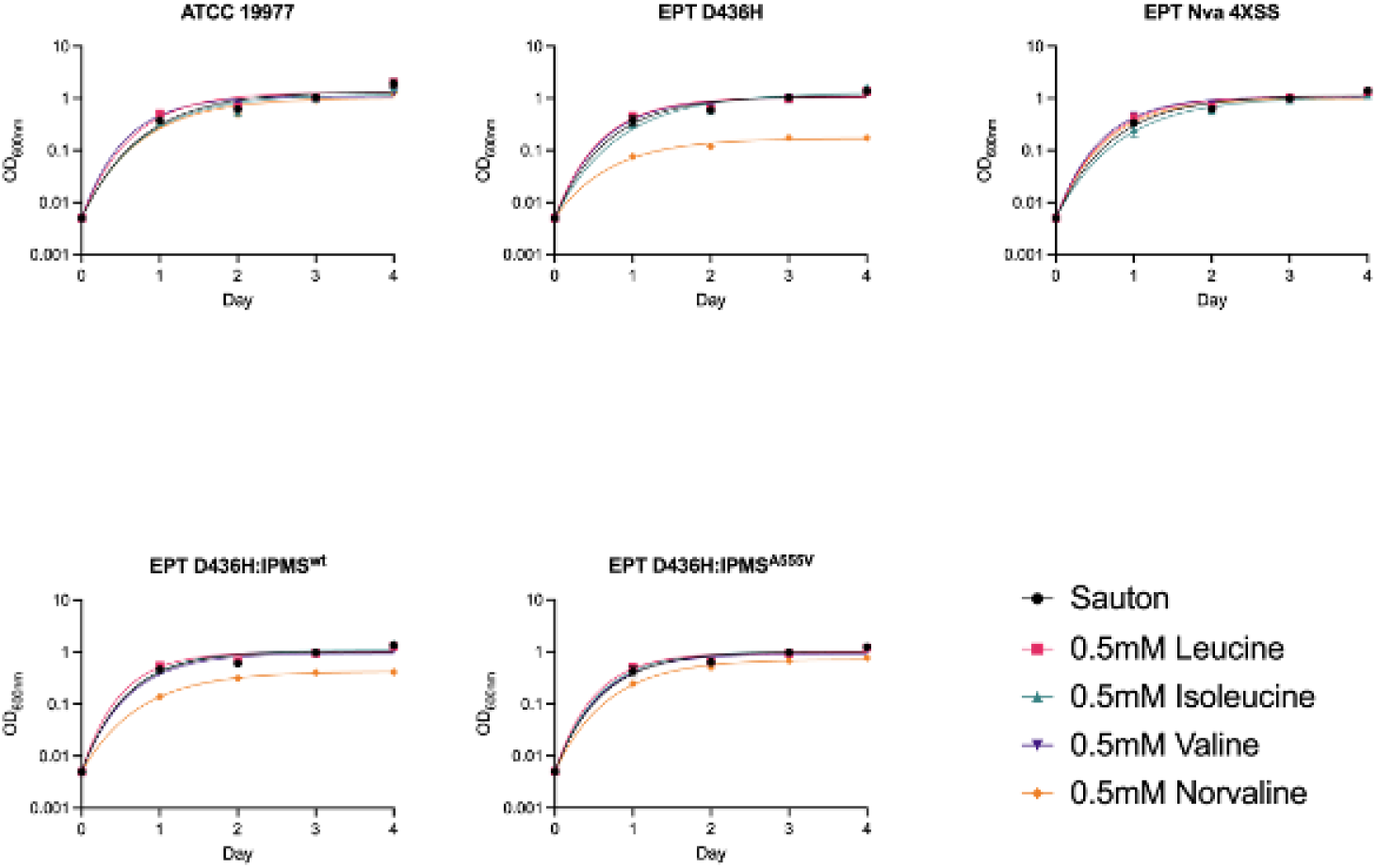
Growth curves of *M. abscessus* complemented with *leuA*. Reference strain ATCC 19977, EPT^R^ mutant LeuRS^D436H^, EPT^R^ mutant complemented with wild type *leuA* or mutant *leuA*, and naturally raised EPT^R^ Nva^R^ mutant were grown statically in Sauton’s minimal media ± l-leucine (pink squares), l-isoleucine (green triangles), l-valine (purple inverted triangles), or l-norvaline (orange diamonds) at 0.5 mM and fit with exponential plateau regression. Data is representative of n = 3 independent experiments with mean ± SD of technical triplicates.

**Fig. S4.**
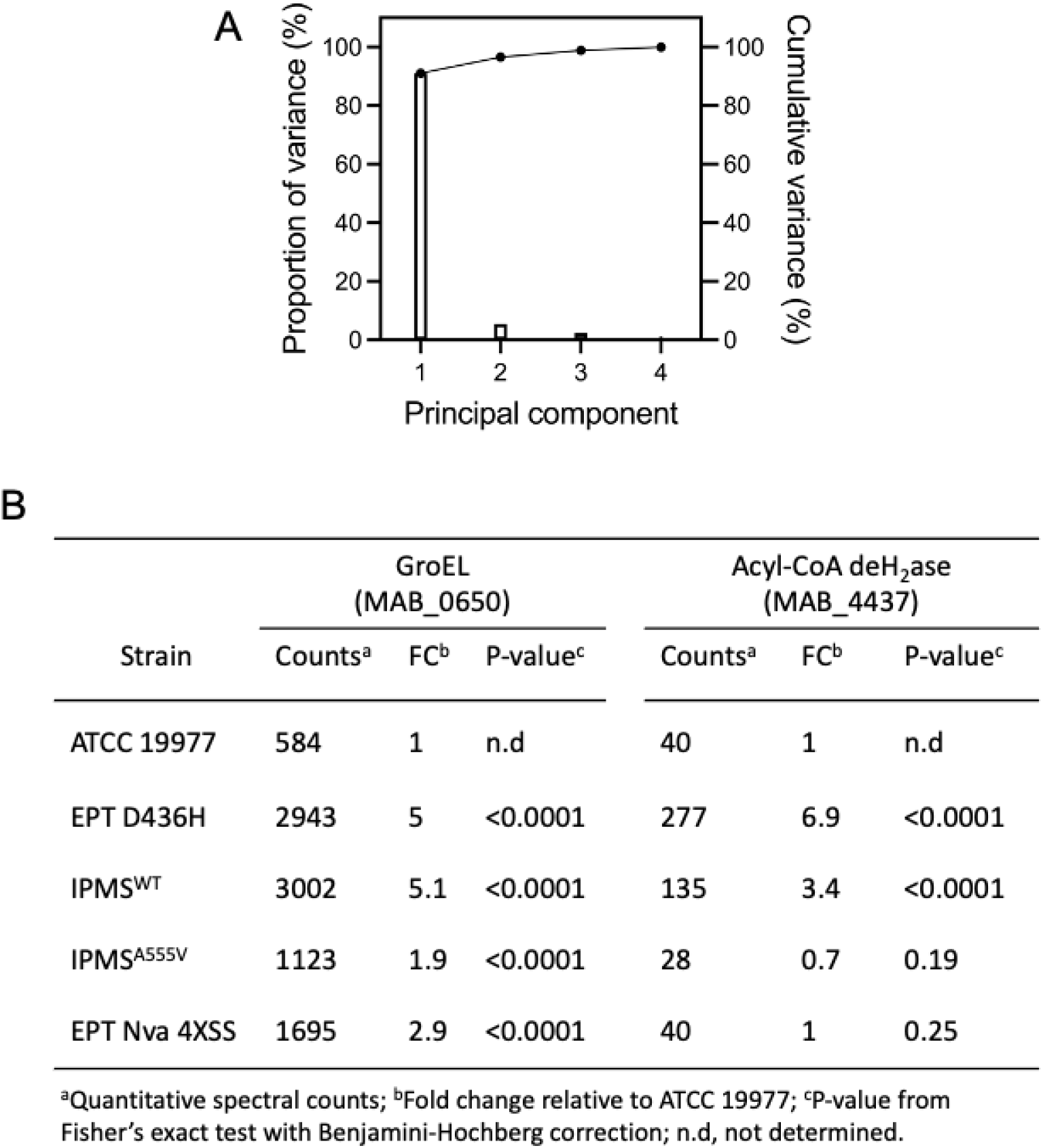
Proteomic analysis of *M. abscessus*. Cell lysates were extracted from *M. abscessus* ATCC 19977, *M. abscessus* EPT^R^ D436H, *M. abscessus* EPT^R^ Nva^R^ 4XSS, *M. abscessus* EPT^R^ D436H complemented with IPMS^WT^ or IPMS^A555V^ grown in Sauton’s media with 0.5 mM l-norvaline for 24 hours. (A) Variance associated with each principal component. PC1 accounted for 90% of the variance, PC2 accounted for 5% of the variance. (B) Quantification of GroEL (largest PC1 coefficient) and Acyl-CoA dehydrogenase (largest PC2 coefficient) abundance.

**Fig. S5.**
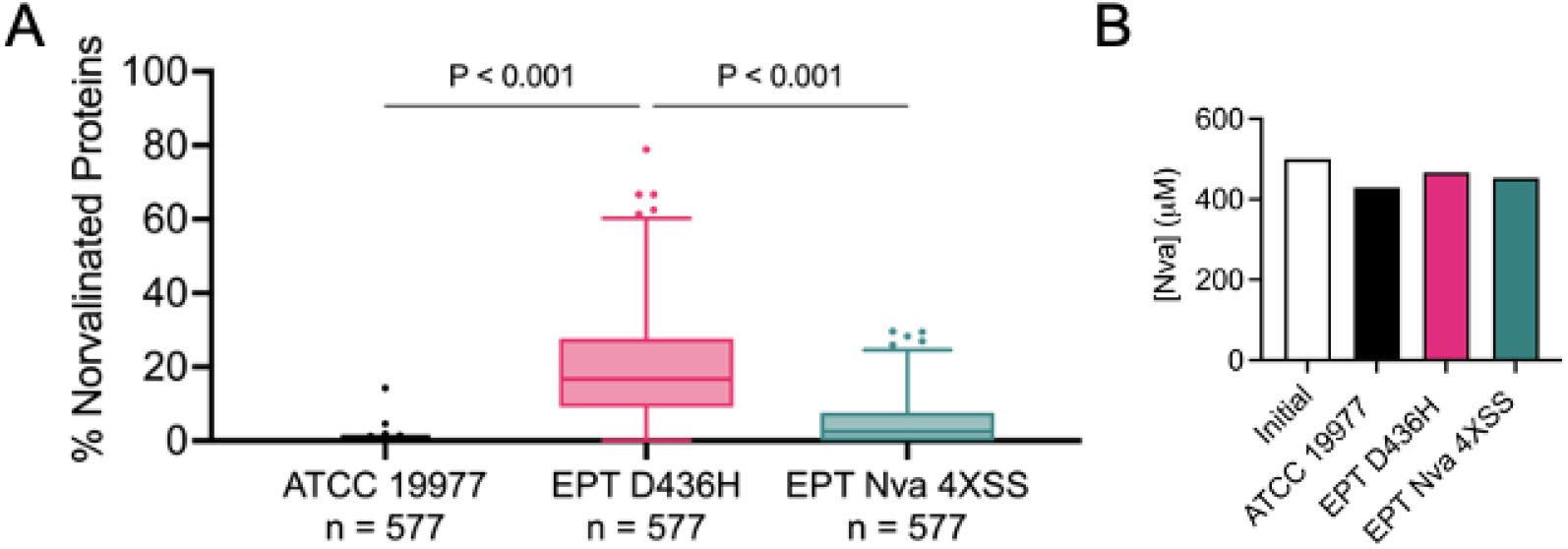
Norvalination of the *M. abscessus* proteome. (A) Percentage of proteins with misincorporation of l-norvaline at leucine residues from cell lysates after 24 h of growth in Sauton’s media with 0.5 mM l-norvaline. Data is median with IQR, whiskers represent 1-99 percentile. P values were obtained by Friedman test with Dunn’s multiple comparisons test. (B) Concentration of l-norvaline remaining in culture supernatants after 24 h l-norvaline challenge from (A).

**Fig. S6.**
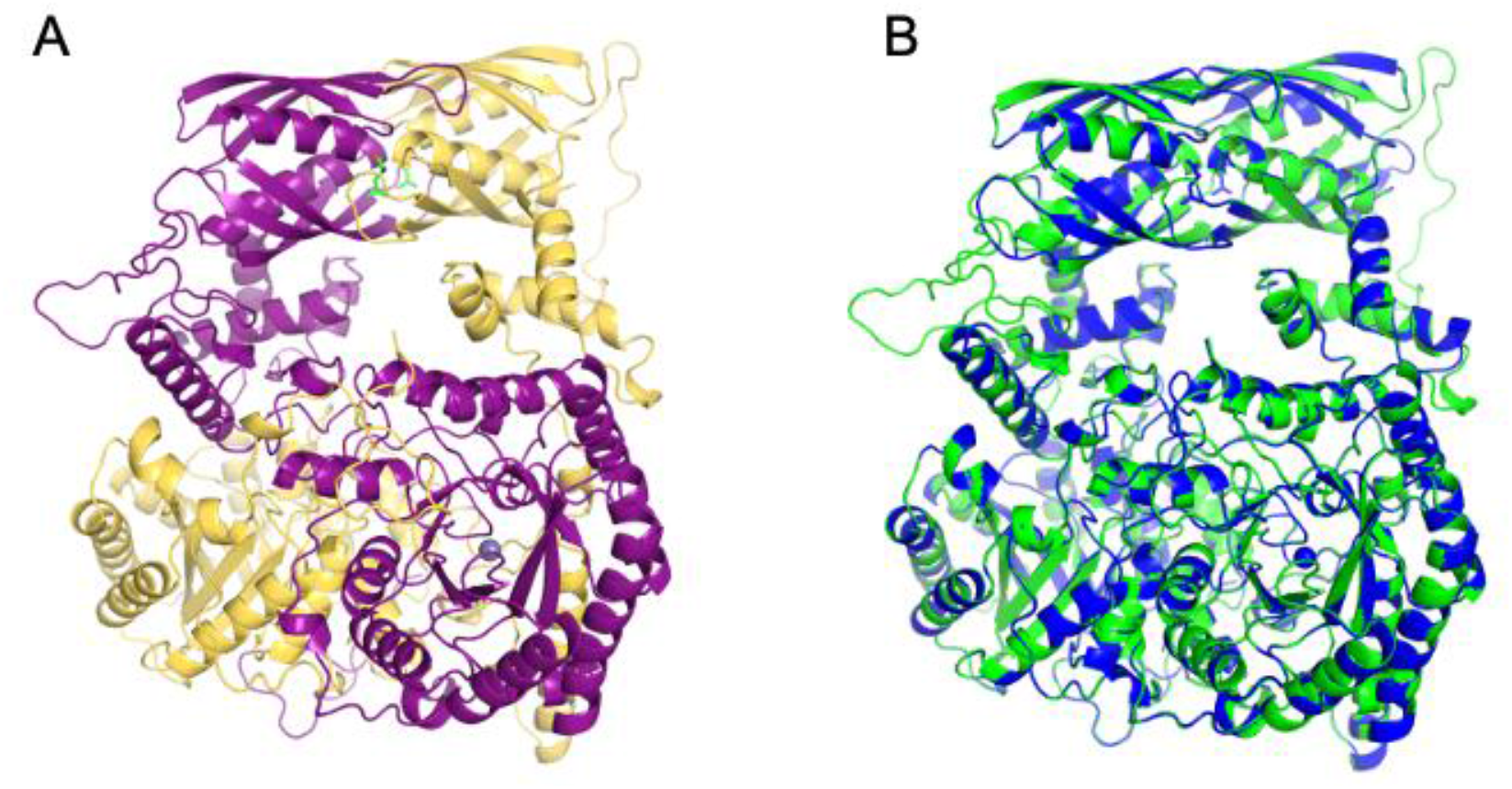
Predicting homology between **α**-IPMS_Mtb_ and **α**-IPMS_Mabs_. (A) **α**-IPMS_Mtb_ model was generated using Swiss-Model and the experimentally determined structure PDB 3FIG with QMEAN global 0.87 ± 0.05. (1–3) (B) Overlay of **α**-IPMS_Mtb_ (green) and Swiss-Model generated **α**-IPMS_Mabs_ (blue) from PDB 3FIG with QMEAN global of 0.86 ± 0.05. Overlay generated rmsd of 0.199 Å.

